# The GATOR complex regulates an essential response to meiotic double-stranded breaks in *Drosophila*

**DOI:** 10.1101/421206

**Authors:** Youheng Wei, Lucia Bettedi, Kuikwon Kim, Chun-Yuan Ting, Mary A. Lilly

**Author notes:** Corresponding author (ML).

## Abstract

The TORC1 inhibitor GATOR1/SEACIT controls meiotic entry and early meiotic events in yeast. However, how metabolic pathways influence meiotic progression in metazoans remains poorly understood. Here we report that the TORC1 regulators GATOR1 and GATOR2 mediate a response to meiotic double-stranded breaks (DSBs) during *Drosophila* oogenesis. We find that meiotic DSBs trigger the activation of a GATOR1 dependent pathway that downregulates TORC1 activity in the female germline. In GATOR1 mutants, high TORC1 activity results in the delayed repair of meiotic DSBs and the hyperactivation of p53. Conversely, the GATOR2 component Mio is required to attenuate GATOR1 activity, to ensure that meiotic DSBs do not trigger a permanent growth arrest. Unexpectedly, we found that GATOR1 inhibits retrotransposon expression in the presence of meiotic DSBs in a pathway that functions in parallel to p53. Our studies have revealed a link between the GATOR complex, the repair of meiotic DSBs and retrotransposon expression

## Introduction

We are interested in understanding how metabolism impacts meiotic progression during oogenesis. <u>T</u>arget <u>o</u>f <u>R</u>apamycin <u>C</u>omplex 1 (TORC1) is a multi-protein complex that functions as a master regulator of metabolism (Jewell & Guan, 2013; Laplante & Sabatini, 2012; Loewith & Hall, 2011). In the presence of adequate nutrients and positive upstream growth signals, TORC1, which contains the serine/threonine kinase Target of Rapamycin, becomes active and functions to stimulate growth and inhibit catabolic metabolism through the phosphorylation of down-stream effector proteins. The <u>Se</u>h1 <u>A</u>ssociated <u>C</u>omplex <u>I</u>nhibits <u>T</u>ORC1 (SEACIT), originally identified in yeast, inhibits TORC1 activity in response to amino acid limitation (Bar-Peled et al., 2013; Dokudovskaya et al., 2011; Neklesa & Davis, 2009; Panchaud, Peli-Gulli, & De Virgilio, 2013; Wu & Tu, 2011). SEACIT, known as the <u>G</u>ap <u>A</u>ctivity <u>To</u>wards <u>R</u>ags complex 1 (GATOR1) in metazoans, is comprised of three highly conserved proteins Npr2/Nprl2, Npr3/Nprl3 and Iml1/Depdc5 (Bar-Peled et al., 2013; Panchaud et al., 2013). In *Drosophila* and mammals, depleting any of the three GATOR1 components results in increased TORC1 activity and growth, as well as a reduced response to amino acid starvation (Bar-Peled et al., 2013; Cai, Wei, Jarnik, Reich, & Lilly, 2016; Dutchak et al., 2015; Kowalczyk et al., 2012; Marsan et al., 2016; Wei et al., 2014). Thus, the role of the SEACIT/GATOR1 complex in the regulation of TORC1 activity is highly conserved in eukaryotes.

The multi-protein GATOR2 complex, known as <u>Se</u>h1 <u>A</u>ssociated <u>C</u>omplex <u>I</u>nhibits <u>T</u>ORC1 (SEACAT) in yeast, inhibits the activity of GATOR1 and thus functions to activate TORC1 (Bar-Peled et al., 2013; Wei et al., 2014). In metazoans, the GATOR2 complex functions in multiple amino acid sensing pathways (Bar-Peled et al., 2013; Cai et al., 2016; Chantranupong et al., 2014; Kim et al., 2015; Panchaud et al., 2013; Parmigiani et al., 2014). In tissue culture cells, depleting GATOR2 components results in the constitutive activation of GATOR1 and the permanent downregulation of TORC1 activity (Bar-Peled et al., 2013; Wei & Lilly, 2014). However, genetic studies of the role of individual GATOR2 components in *Drosophila*, indicate that the requirement for the GATOR2 complex is more nuanced when examined in the context of a multicellular animal. For example, mutations in the GATOR2 component *mio*, result in a block to oocyte growth and differentiation, due to the constitutive activation of the TORC1 inhibitor GATOR1 in the female germline (Iida & Lilly, 2004; Wei, Reveal, Cai, & Lilly, 2016). However, *mio* is not required to maintain TORC1 activity in most somatic tissues of *Drosophila* (Wei et al., 2016). Why there is a tissue specific requirement for *mio* in the female germline of *Drosophila* is currently unknown.

In single celled eukaryotes, nutrient limitation often facilitates meiotic entry (van Werven and Amon, 2011). In the yeast *Saccharomyces cerevisiae*, the down-regulation of TORC1 by SEACIT/GATOR1 in response to amino acid stress promotes both meiotic entry and early meiotic progression (Deutschbauer, Williams, Chu, & Davis, 2002; Jordan, Klein, & Leach, 2007; Neklesa & Davis, 2009; Spielewoy et al., 2010). Surprisingly, as is observed in yeast, during *Drosophila* oogenesis the GATOR1 complex promotes meiotic entry (Wei et al., 2014). These data raise the intriguing possibility that in *Drosophila* the GATOR1 complex and low TORC1 activity may be critical to the regulation of additional events of the early meiotic cycle.

Here we report that the GATOR complex regulates an essential response to meiotic DSB during *Drosophila* oogenesis. We demonstrate that in the female germline of *Drosophila*, meiotic DSBs trigger a GATOR1 dependent downregulation of TORC1 activity which promotes the timely repair of meiotic DSBs and prevents the hyperactivation of p53. Conversely, the GATOR2 component Mio opposes the activity of GATOR1 in the female germline, thus allowing for the growth and development of the oocyte in later stages of oogenesis. Thus, we have identified a GATOR complex based regulatory loop that modulates TORC1 activity in response to meiotic DSBs. Additionally, during the course of our studies, we observed that the GATOR1 complex prevents the derepression of retrotransposon in the presence of meiotic DSBs.

## Results

### Mio attenuates a GATOR1 dependent response to meiotic DSBs

The GATOR2 complex inhibits the TORC1 inhibitor GATOR1 (Fig 1A). Ovaries from mutants of the GATOR2 component *mio*, have reduced TORC1 activity and are severely growth restricted (Fig. 1B-D) (Iida & Lilly, 2004; Wei et al., 2014). In our previous studies, we demonstrated that the *mio* ovarian phenotypes result from the constitutive downregulation of TORC1 activity via a GATOR1 dependent pathway (Wei et al., 2014). Thus, removing GATOR1 activity in the *mio* mutant background, as is observed in *mio, nprl3* double mutants, results in increased TORC1 activity and rescues the *mio* ovarian phenotypes (Wei et al., 2014).

**Figure 1.**
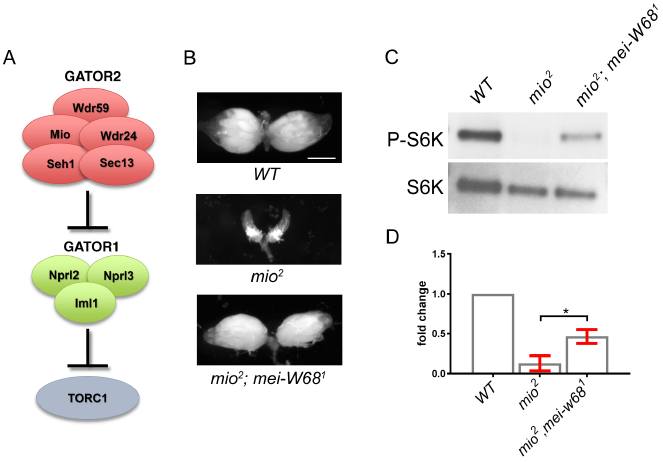
Meiotic DSBs trigger the downregulation of TORC1 activity during *Drosophila* oogenesis. **(A)** The GATOR2 complex opposes the activity of the TORC1 inhibitor GATOR1. **(B)** Representative ovaries from wild type (WT), *mio*^*2*^ and *mio2, mei-W68*^*1*^ females. Scale bar, 30 µm. **(C)** Western blot of phospho-S6K and total-S6K levels of whole ovaries prepared from WT, *mio*^*2*^ and *mio*^*2*^, *mei-w68*^*1*^ mutant females. **(D)** Quantification of phospho-S6K levels relative to the total S6K. Error bars represent the standard deviation for three independent experiments. Unpaired T-student test was used to calculate the statistical significance. * P <0.05.

Surprisingly, the *mio* mutants are also suppressed by blocking the formation of meiotic DSBs, with approximately 70% of *mio* ovaries achieving wild-type levels of growth when double mutant for genes required to generate meiotic DSBs (Fig. 1B) (Iida & Lilly, 2004). One model to explain this observation is that meiotic DSBs promote the GATOR1 dependent downregulation of TORC1 activity in the early meiotic cycle and that *mio* is required to oppose or attenuate this response. To test this idea, we examined if blocking the formation of meiotic DSBs in the *mio* mutant background resulted in increased TORC1 activity. Towards this end, we compared the phosphorylation status of S6 kinase, a downstream TORC1 target, in ovaries from *mio* single mutants versus *mio, mei-W68* double mutant ovaries (Fig 1C,D). *mei-W68* (*SPO11* homolog) is a highly-conserved enzyme required for the generation of meiotic DSBs (McKim and Hayashi-Hagihara, 1998; Sekelsky et al., 1999; Liu et al., 2002). Notably, we found that relative to ovaries from *mio* single mutants, *mio, mei-W68* double mutants exhibit a dramatic increase in TORC1 activity (Fig 1C,D). Thus, the GATOR1 dependent pathway that triggers the constitutive downregulation of TORC1 activity in *mio* mutants is activated, at least in part, by the presence of meiotic DSBs.

### GATOR1 promotes the repair of meiotic DSB

In previous work we demonstrated that the GATOR1 complex downregulates TORC1 activity to facilitate meiotic entry in *Drosophila* ovarian cysts (Deutschbauer et al. 2002; Jordan et al. 2007; Neklesa and Davis 2009; Spielewoy et al. 2010; Wei et al. 2014). However, the delay in meiotic entry observed in *Drosophila* GATOR1 mutants is not fully penetrant and therefore unlikely to be the sole cause of the infertility observed in mutant females (Fig. S1) (Wei et al. 2014). Considering our findings that meiotic DSBs serve to promote and/or reinforce low TORC1 activity after the mitotic/meiotic switch, we hypothesized that as is observed in yeast, the GATOR1 dependent downregulation of TORC1 activity may be critical to the regulation of additional early meiotic events, including the repair of meiotic DSBs in *Drosophila*.

To test this hypothesis, we examined the behavior of meiotic DSBs in GATOR1 mutant oocytes. The programmed generation of DSBs by the Spo11/Mei-W68 enzyme initiates meiotic recombination in early prophase of meiosis I (Hunter, 2015; McKim & Hayashi-Hagihara, 1998). During *Drosophila* oogenesis, the kinetics of DSB formation and repair can be followed using an antibody against the phosphorylated form of His2Av known as γ-His2Av (Madigan, Chotkowski, & Glaser, 2002; Mehrotra & McKim, 2006). *Drosophila* ovarian cysts generate meiotic DSBs after the initiation of synaptonemal complex (SC) formation in region 2a of the germarium (Fig. 2A) (Carpenter, 1975; Jang, Sherizen, Bhagat, Manheim, & McKim, 2003; Mehrotra & McKim, 2006). γ-H2Av nuclear foci are first observed in the two pro-oocytes, which are in early pachytene (Jang et al., 2003; Mehrotra & McKim, 2006). Subsequently, a small number of DSBs are also observed in the pro-nurse cells (Mehrotra & McKim, 2006; Narbonne-Reveau & Lilly, 2009). As meiosis proceeds and the DSBs are repaired, γ-H2Av-positive foci decrease in number and mostly disappear by late region 2b (Jang et al., 2003; Mehrotra & McKim, 2006). γ-H2Av signals are rarely detected in region 3 oocytes (Fig. 2A, arrow). Analysis of mutants that fail to repair DSBs, and thus capture the total number of meiotic breaks, indicate that wild-type oocytes generate approximately 20-25 Spo11/Mei-W68 dependent breaks during oogenesis (Joyce et al., 2011; Mehrotra & McKim, 2006).

**Figure 2.**
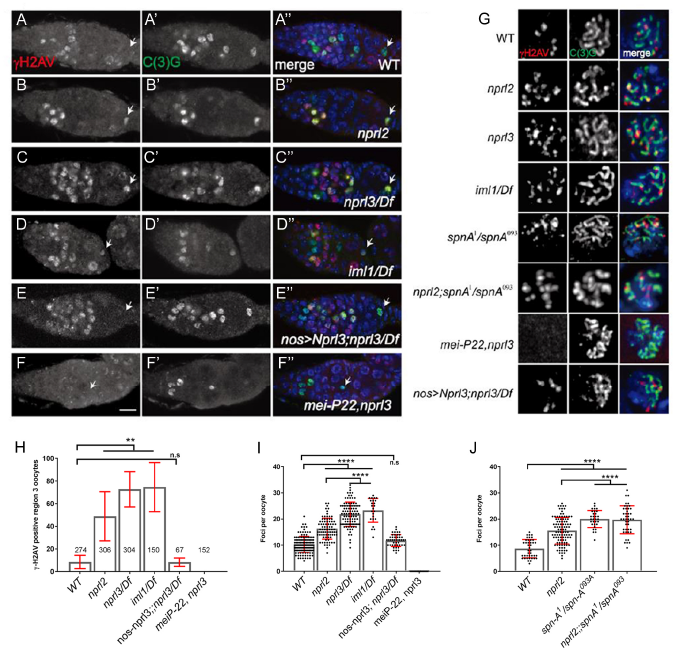
GATOR1 influences the number and persistence of DSBs in early oocytes. Ovaries from **(A)** wild type, **(B)** *nprl2*^*1*^, **(C)** *nprl3*^*1*^*/Df*, **(D)** *iml*^*1*^*/Df*, **(E)** *nanos-GAL4; UAS-Nprl3; nprl3*^*1*^*/Df* and **(F)** *mei-P22*^*P22*^, *nprl3*^*1*^ flies were stained for C(3)G (green, **A’-F’**) and γ - H2Av (red, **A-F**). C(3)G marks the synaptonemal complex (SC) and is used to mark oocytes and follow meiotic progression. γ-H2Av marks DSBs. Scale bars, 10µm. In wild type oocytes, meiotic DSBs are induced in region 2a and repaired by region 3 (arrow). In GATOR1 mutants, DSBs persist in region 3 oocytes. In *nanos-GAL4; UAS-Nprl3; nprl3*^*1*^*/Df* oocytes, DSBs are repaired by region 3. In *mei-P22*^*P22*^, *nprl3*^*1*^ mutants have no DSBs. **(G)** Ovaries from wild type, *nprl2*^*1*^, *nprl3*^*1*^*/Df, iml*^*1*^*/Df, spn-A*^*1*^*/spn-A*^*093A*^, *nprl2*^*1*^; *spnA*^*1*^*/spnA*^*093*^,*mei-P22*^*p22*^,*nprl3*^*1*^ and *nanos-GAL4; UAS-Nprl3; nprl3*^*1*^*/Df* flies were stained for C(3)G (green) and γ-H2Av (red). Representative immunofluorescent microphotographs of the γ-H2Av foci in region 2a oocyte are shown. **(H)** Percentage of region 3 oocytes with γ-H2Av foci. **(I and J)** Quantification of γ-H2Av foci in region 2a oocytes. Error bars represent SD from at least three independent experiments.** P<0.001, ***P < 0.001, ns: no significance.

To determine if GATOR1 regulates the behavior of meiotic DSBs in *Drosophila*, we compared the pattern of γ-H2Av foci in wild-type versus GATOR1 mutant ovaries with antibodies against γ-H2Av and the SC component C(3)G, to highlight DSBs and oocytes respectively (Iida & Lilly, 2004; Mehrotra & McKim, 2006). We determined that while the majority of wild-type oocytes had repaired all of their DSBs and thus had no γ-H2Av foci by region 3 of the germarium, in GATOR1 mutants between 50%-80% of region 3 oocytes are γ-H2Av positive (Fig. 2A-E, H, arrow) (Mehrotra & McKim, 2006). Moreover, GATOR1 mutant oocytes had a significant increase in the steady state number of γ-H2Av foci per oocyte nucleus in region 2A of the germarium relative to wild-type oocytes (Fig. 2G). From these data, we conclude that the GATOR1 complex influences the behavior of meiotic DSB during early oogenesis.

Next, we examined if the altered γ-H2Av pattern observed in GATOR1 mutants was dependent on the meiotic DSB machinery. Alternatively, the extra DSB may be induced during the premeiotic S phase, as is observed in mutants of the CdkE/Cdk2 inhibitor *dacapo* (Hong, Lee-Kong, Iida, Sugimura, & Lilly, 2003). To address this question, we generated double-mutants containing a null allele for *mei-p22*, a gene required for the formation of DSBs in *Drosophila* (H. Liu, Jang, Kato, & McKim, 2002) and a null allele for the GATOR1 component *nprl3* (Cai et al., 2016). We determined that double-mutant *nprl3, mei-p22* oocytes had no *γ*-H2Av foci (Fig. 2G,I). These data indicate that the increase in the steady state number of γ-H2Av foci, as well as the retention of these foci into region 3 of the germarium, are dependent on the meiotic DSB machinery.

During the development of wild-type *Drosophila* oocytes, asynchrony in the generation and repair of meiotic DSBs ensures that the number of γ-H2Av foci observed in an oocyte at any single time point, is less than the total number of DSBs generated during the lifetime of the oocyte (Joyce et al., 2011; Mehrotra & McKim, 2006). We noticed that the number of γ-H2Av foci observed in GATOR1 mutant oocytes is never more than the 20-25 foci observed of mutants in the DSB repair pathway (Fig. 2I) (Mehrotra & McKim, 2006). This observation suggested that the increase in the steady state number of DSBs observed in GATOR1 mutants may result from the delayed repair of meiotic DSBs, rather than the production of extra Mei-W68/Spo11 induced DSBs. To test this hypothesis, we generated *spnA*^*1*^*/spnA*^*093A*^*nprl2*^*1*^ double mutants. Importantly, *spnA*^*1*^*/spnA*^*093A*^transherterozyogtes fail to repair meiotic DSBs (Joyce & McKim, 2011; Staeva-Vieira, Yoo, & Lehmann, 2003). If *nprl2* mutants make extra meiotic breaks, then the number of foci in the *nprl2, spnA* double mutant should be higher than either single mutant. However, we found that the *spnA*^*1*^*/spnA*^*093*^ *nprl2*^*1*^ double-mutants contained approximately the same number of γ-H2Av foci as *spnA*^*1*^*/spnA*^*093*^ single mutants (Fig. 2J). Thus, mutations in *nprl2* do not result in the production of extra meiotic breaks. Taken together, our data strongly suggest that the GATOR1 complex influences the repair, rather than the production, of meiotic DSBs.

### GATOR1 mutants hyperactivate p53 in response to meiotic DSBs

p53, a transcription factor that mediates a conserved response to genotoxic stress, regulates early meiotic events in multiple organisms (S. Lee, Cavallo, & Griffith, 1997; Linke et al., 2003; Mateo et al., 2016; Sturzbecher, Donzelmann, Henning, Knippschild, & Buchhop, 1996). During *Drosophila* oogenesis, the generation of meiotic DSBs results in the brief activation of p53 and the expression of downstream targets (Fig. 3A) (Lu, Chapo, Roig, & Abrams, 2010). To determine if GATOR1 mutants experience increased genotoxic stress during oogenesis, we used a reporter construct to assay p53 activity. The p53-GFPnls reporter construct contains the Green Fluorescent Protein (GFP) under the control of an enhancer from the p53 transcriptional target *reaper* (Lu et al., 2010). In wild-type ovaries, a faint signal from the p53-GFPnls reporter is first observed in region 2a of the germarium, concurrent with the generation of meiotic DSBs (Fig. 3A, arrow) (Lu et al., 2010). As ovarian cysts continue to develop, the p53-GFPnls signal rapidly dissipates as meiotic DSBs are repaired (Fig. 3A, arrowhead) (Lu et al., 2010). By region 3 of the germarium less than 5% of p53-GFPnls ovarian cysts contain detectable levels of GFP (Fig. 3B). In contrast, the germaria of all three GATOR1 mutants exhibited a dramatic increase in both the strength and the duration of p53-GFPnls expression in the germarium, with strong GFP signal observed in nearly 80% of region 3 ovarian cysts (Fig. 3A-D, F, G, arrowheads). Homozygous germline clones of the *iml1*^*1*^ and *nprl3*^*1*^ null alleles hyperactivate p53 confirming that GATOR1 activity is required cell autonomously in the female germline (Fig. S2).

**Figure 3.**
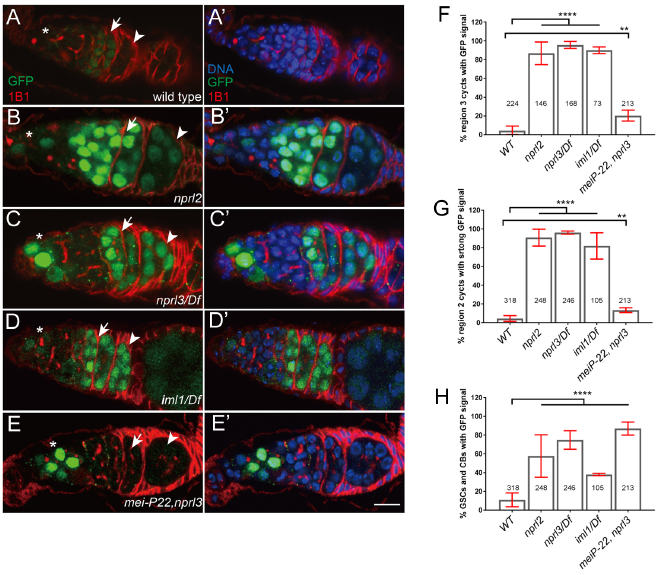
GATOR1 prevents p53 hyperactivation in Drosophila early ovarian cysts. Ovaries from **(A)** *p53R-GFP*, **(B)** *nprl2*^*1*^; *p53R-GFP*, **(C)** *p53R-GFP;nprl3*^*1*^*/Df*, **(D)** *p53R-GFP; iml*^*1*^*/Df* and **(E)** *p53R-GFP; mei-P22*^*p22*^, *nprl3*^*1*^ were stained for GFP (green) and 1B1 (red). Germarial regions are defined by *1B1* staining. In wild-type ovaries the p53-GFP reporter is briefly activated in region 2 (indicated by arrow). Note the low level of GFP staining. In contrast, in GATOR1 mutants, p53R-GFP is robustly activated with strong GFP signal often persisting into germarial region 3 and beyond. Additionally, in GATOR1 mutant germaria, p53R-GFP is frequently activated in germline stem cell (GSC) and daughter cystoblasts (CB). In *mei-P22*^*p22*^, *nprl3*^*1*^ double mutant germaria, the hyperactivation *of* p53R-GFP is rescued in region 2a ovarian cysts. However, p53-GFP activation in GSC and CB is retained in the double mutants (asterisk) indicating that in these cells the activation of p53 is not contingent on the presence of meiotic DSBs. Scale bars, 10µm. **(F)** Percentage of germaria with sustain p53R-GFP signal in region 3. **(G)** Percentage of germaria with high p53R-GFP signal in region 2. **(H)** Percentage of germaria with p53R-GFP expression in GSC and CB. Error bars represent SD from at least three independent experiments. **P < 0.01, ****P<0.0001.

We predicted that the persistent hyperactivation of p53 in the female germ line of GATOR1 mutants is due to the delayed repair of meiotic DSBs. To test this model, we examined p53 activation in the *mei-p22, nprl3* double mutant. Strikingly, inhibiting meiotic DSBs strongly suppressed the expression of p53-GFPnls in the *nprl3* mutant background (Fig. 3E). From these observations, we infer that in GATOR1 mutant ovaries, the hyperactivation of p53 is downstream of Spo11/Mei-W68-induced DBS. Moreover, we conclude that the GATOR1 complex is required to oppose genotoxic stress triggered by meiotic DSBs and/or other downstream events of meiotic recombination.

Recent evidence indicates that p53 is activated in germline stem cells of *Drosophila* after exposure to cellular stresses, including deregulated growth and ionizing radiation (Wylie et al. 2014; Ma et al. 2016). The GATOR1 complex inhibits TORC1 activity and is required to restrain growth in *Drosophila* (Wei et al. 2014; Cai et al. 2016). As was reported with other mutants that deregulate growth (Wylie et al. 2014), we found that the p53-GFPnls reporter construct is robustly activated in the germline stem cells and their near descendants in GATOR1 mutant females (Fig. 3B-D, H). Consistent with the restriction of Mei-W68/Spo11 activity to meiotic cysts, the *mei-p22, nprl3* double mutants retain p53-GFPnls expression in stem cells even though the meiotic activation of the p53-GFPnls reporter is lost in regions 2a and 2b of the germarium in the *mei-p22, nprl3* double mutant (Fig. 3E-G). Thus, an independent cellular stress, likely related to deregulated metabolism or growth, activates p53 in the germline stem cells and cystoblasts of GATOR1 mutant females.

### GATOR1 mutant larvae are sensitive to genotoxic stress

The <u>T</u>uberous <u>s</u>clerosis <u>c</u>omplex (TSC) is a potent inhibitor of TORC1 that directly inhibits the small GTPase Rheb, a critical activator of TORC1 (Inoki, Li, Xu, & Guan, 2003; Zhang et al., 2003). In humans, mutations in components of TSC sensitize cells to multiple forms of genotoxic stress (Deschavanne & Fertil, 1996; C. H. Lee et al., 2007; Pai et al., 2016; Paterson, Sell, Smith, & Bech-Hansen, 1982). Thus, we predicted that the GATOR1 mutants, which have a two to three-fold increase in TORC1 activity, might have a globally diminished ability to respond to DNA damage. To test this model, we treated *nprl2* and *nprl3* mutant larvae with the mutagen Methyl Methane Sulfonate (MMS) and compared the percentage of mutant animals that survived to adulthood relative to sibling heterozygous controls. MMS generates an array of DNA lesions including DSBs (Magana-Schwencke, Henriques, Chanet, & Moustacchi, 1982). Notably, we found that GATOR1 mutant larvae were sensitive to DNA damage, with *nprl3/Df* transheterozygotes exhibiting a greater than 10-fold decrease in survival rates when exposed to 0.08% MMS (Table 1). These data support the idea that the GATOR1 complex plays a critical role in the response to genotoxic stress in germline and somatic tissues.

**Table 1.** *nprl2* and *nprl3* larvae are sensitive to the mutagen methyl methane sulfonate

| **Genotype** | **0%MMS** | **0.04% MMS** | **0.08% MMS** | **n** |
| --- | --- | --- | --- | --- |
| *nprl2*<sup>1</sup>/*nprl2*<sup>1</sup>(%) | 29 (2.4) | 13 (5.2) | 7 (4.1) | 624, 561, 307 |
| *nprl3*<sup>1</sup>/*Df*(%) | 27 (0.1) | 8 (3.7) | 2 (1.1) | 1537, 1171, 904 |

Percentages are shown as the fraction of the total homozygous mutants among all surviving progeny. The larvae treated with MMS were derived from heterozygous parents and the surviving adult progeny were determined. *nprl2* was balanced over FM7a while *nprl3* and *Df(3L)ED4515, P{3'.RS5+3.3'}ED4515* were balanced over TM3, Sb. In parenthesis are the standard errors between experiments and n indicates the total number of progeny counted in all independent experiments. Mutant progeny were expected to constitute 33.3% of the progeny. Note that for *nprl2*, which is on the X chromosome and thus hemizygous in males, the data reflect the percentage of homozygous female progeny among all female progeny.

### *Tsc1* germline depletions phenocopy GATOR1 mutants

The most parsimonious interpretation of our data is that in GATOR1 mutant ovaries, high TORC1 activity opposes the timely repair of meiotic DSBs and increases genotoxic stress. To test this model, we depleted the TSC component Tsc1 from the female germline. As described above TSC is a potent inhibitor of TORC1 that directly inhibits the small GTPase Rheb, a critical activator of TORC1 (Inoki et al., 2003; Zhang et al., 2003). We determined that depleting *Tsc1* in the female germline using RNAi, resulted in the robust expression of p53-GFPnls during the early meiotic cycle (Fig. 4A-C). Moreover, as was observed in GATOR1 mutants, *Tsc1*^*RNAi*^ oocytes had an increase in the steady state number of γ-H2Av foci as well as an increase in the percentage of oocytes that retained γ-H2Av positive into region 3 of the germarium (Fig. 4E, F). Taken together our data support the model that the tight control of TORC1 activity by both GATOR1 and TSC is essential to the proper regulation of meiotic DSBs during *Drosophila* oogenesis.

**Figure 4.**
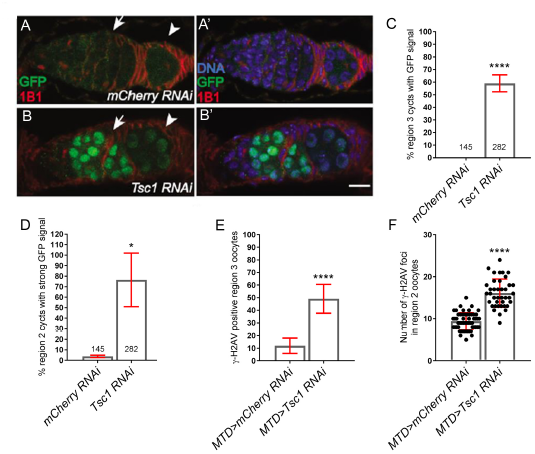
The TORC1 inhibitor TSC1 promotes genomic stability in early oocytes. Ovaries from **(A)** *p53R-GFP; MTD>mCherry RNAi* and **(B)** *p53R-GFP; MTD>Tsc1 RNAi* flies were stained for GFP (green) and 1B1 (red). In the *mCherry RNAi* (control) ovaries the p53-GFP expression is very low. In contrast, the *Tsc1 RNAi* ovaries with strong GFP signal and often persisting into germarial region 3. Scale bars, 10µm. **(C)** Percentage of germaria with strong p53R-GFP signal in region 3. **(D)** Percentage of germaria with sustained p53R-GFP signal in region 3. The γ-H2Av foci were determined in *MTD>mCherry RNAi* and *MTD>Tsc1 RNAi* ovaries. **(E)** Percentage of region 3 oocytes with γ-H2Av foci. **(F)** Quantification of γ-H2Av foci per oocyte in region 2a. Error bars represent SD from at least three independent experiments. *P < 0.05, ****P<0.0001.

### GATOR1 inhibits retrotransposon expression in *Drosophila*

Recent evidence implicates p53 in the repression of retrotransposon activation (Levine, Ting, & Greenbaum, 2016; Wylie et al., 2016). Intriguingly, in the female germ line of *Drosophila*, p53 is only required to inhibit retrotransposon expression in the presence of meiotic DSBs (Wylie et al., 2016). These results are consistent with the model that genotoxic stress promotes TE activation, although a direct role for p53 in the inhibition of retrotransposon expression has also been proposed (Harris et al., 2009; McClintock, 1984; Wylie et al., 2016). We wanted to know if retrotransposons expression is similarly derepressed in the ovaries of GATOR1 mutants. Towards this end, we used qRT-PCR to compare expression levels for multiple retrotransposons in wild type versus *nprl2* and *nprl3* mutant ovaries. We found that *nprl2* and *nprl3* mutant ovaries have increased expression of multiple retrotransposons including TAHRE, Het-A, Indefix and Gypsy (Fig. 5A). In contrast to *nprl2* and *nprl3* mutants, *Tsc1*^*RNAi*^ germline resulted in little or no increase in retrotransposon expression (Fig. S3). From these results, we conclude that the GATOR1 components *nprl2* and *nprl3* oppose retrotransposon expression in the female germline of *Drosophila*.

**Figure 5.**
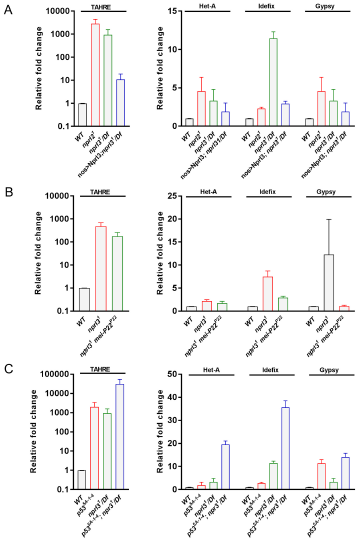
GATOR1 opposes retrotransposon expression in parallel to p53. **(A)** Quantitative RT-PCR analysis of expression levels for retrotransposons in wild type, *nprl2*^*1*^, *nprl3*^*1*^*/Df* and *nanos-GAL4; UAS-Nprl3; nprl3*^*1*^*/Df* ovaries. **(B)** Quantitative RT-PCR analysis of expression levels for transposons in wild type, *nprl3*^*1*^ and *mei-P22*^*P22*^, *nprl3*^*1*^ ovaries. **(C)** Quantitative RT-PCR analysis of expression levels for the transposons in wild type, *p53*^*5*^A-^1-4^, *nprl3*^*1*^*/Df* and *nprl3*^*1*^*/Df, p53*^*5A-1-4*^ ovaries. Rp49 is used for normalization. Fold expression levels are relative to wild type.

In *nprl3* mutant ovaries, the activation of p53 and the increase in *γ*-H2Av foci is dependent on the production of meiotic DSBs (Fig. 2 and 3). To determine if the expression of retrotransposons in *nprl3* mutant ovaries also requires the meiotic DSB machinery, we examined *nprl3, mei-p22* double mutants. Using qRT-PCR we observed that retrotransposon expression was largely, but not completely, suppressed in *nprl3, mei-p22* double-mutant ovaries. Thus, meiotic DSBs trigger the expression of retrotransposons during *Drosophila* oogenesis in the *nprl3* mutant background (Fig. 5B). However, it is important to note that these data suggest that the GATOR1 complex may also impact retrotransposon expression independent of meiotic DSBs as indicated by the relatively modest rescue of TAHRE over-expression observed in *nprl3, mei-p22* double mutants (Fig. 5B).

Finally, we wanted to determine if the GATOR1 complex inhibits the activation of retrotransposons through the p53 pathway. To answer this question, we performed epistasis analysis by generating double mutants that were homozygous for null alleles of both *p53* and *nprl3*. Strikingly, the *p53, nprl3* double mutant ovaries showed a dramatic increase in retrotransposon expression relative to either single mutant (Fig. 5C). Thus, the *p53* and *nprl3* phenotypes are additive with respect to the inhibition of retrotransposon expression. These data strongly suggest that GATOR1 and p53 act through independent pathways to inhibit retrotransposon activation in the female germline during meiosis (Fig. 5C).

## Discussion

Recent evidence implicates metabolic pathways as important regulators of meiotic progression and gametogenesis (Chi et al., 2016; Ferguson, Blundon, Klovstad, & Schupbach, 2012; Guo et al., 2018; LaFever, Feoktistov, Hsu, & Drummond-Barbosa, 2010; Sieber, Thomsen, & Spradling, 2016). Here we define a role for the GATOR complex, a conserved regulator of TORC1 activity, in the regulation of two events that impact germline genome stability: the response to meiotic DSBs and the inhibition of retrotransposon expression.

### GATOR2 opposes a GATOR1 dependent response to meiotic DSBs

We have previously shown that in *Drosophila*, mutations in the GATOR2 component *mio*, result in the constitutive activation of the GATOR1 pathway in the female germline but not in somatic tissues (Iida & Lilly, 2004; Wei et al., 2014). Here we demonstrate that the tissue specific requirement for Mio during oogenesis is due, at least in part, to the generation of meiotic DBSs during oogenesis. In *Drosophila*, only the female germline undergoes meiotic recombination and thus experiences the genotoxic stress associated with developmentally programmed DSBs (Hughes, Miller, Miller, & Hawley, 2018). We find that in *mio* mutants, blocking the formation of meiotic DSBs prevents the constitutive downregulation of TORC1 activity thus allowing for the growth and development of the oocyte. These data are consistent with the model that meiotic DSBs trigger the activation of a GATOR1 dependent pathway that must be opposed and/or attenuated by the GATOR2 component Mio.

There are at least two possible models, which are not mutually exclusive, for how meiotic DSBs impact TORC1 regulation (Figure 6). First, the GATOR2 complex may retain the ability to downregulate GATOR1 activity even in the absence of the Mio subunit. In this model, meiotic DSBs send a signal that fully inactivates the GATOR2 dependent inhibition of GATOR1. However, we favor a second model in which meiotic DSBs result in the activation of the TORC1 inhibitor TSC. We previously reported that, as is observed with GATOR1, depleting TSC complex components in the *mio* mutant background rescues the *mio* ovarian phenotypes (Wei et al., 2014). Moreover, we determined that when TSC is constitutively active in the female germline, as is observed upon the depletion of the TSC inhibitor Akt1, ovarian growth is dramatically reduced (Wei & Lilly, 2014). Consistent with the similar of GATOR1 and TSC depletions in the female germline, recent reports indicate that GATOR1 and TSC can act in a common pathway to affect the full downregulation of TORC1 activity in response to multiple upstream inhibitory inputs (Demetriades, Doumpas, & Teleman, 2014; Demetriades, Plescher, & Teleman, 2016; Menon et al., 2014). The determination of the exact mechanism by which meiotic DSBs trigger the activation of the GATOR1 and/or TSC pathway will be an exciting area for future studies.

**Figure 6.**
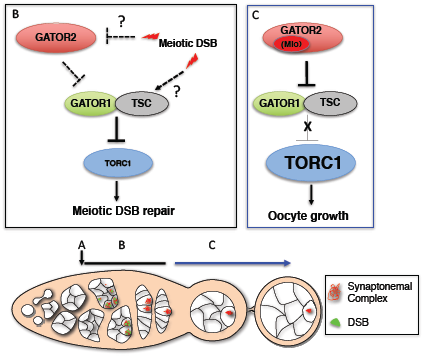
The GATOR complex regulates a response to meiotic DSBs during *Drosophila* oogenesis. **(A)** The downregulation of TORC1 activity by the GATOR1 complex promotes meiotic entry (Wei, et al., 2014). **(B)** After meiotic entry, meiotic DSBs activate a GATOR1/TSC dependent pathway to ensure low TORC1 activity in early prophase of meiosis I. This may occur either through the inhibition of the GATOR2 complex or the activation of the TORC1 inhibitor TSC. See text for details. Low TORC1 activity promotes the timely repair of meiotic DSBs. **(C)** Subsequently, the GATOR2 component Mio, attenuates the activity of the GATOR1/TSC pathway, thus allowing for increased TORC1 activity and the growth and development of the oocyte in later stages of oogenesis.

The inhibition of meiotic DSBs results in a substantial, but not complete, suppression of the *mio* ovarian phenotypes with respect to growth and TORC1 activity (Iida and Lilly 2004). Previously, we demonstrated that a GATOR1 dependent pathway promotes meiotic entry early in oogenesis, prior to the generation of meiotic DSBs (Wei et al. 2014). Thus, the inability to fully suppress the *mio* phenotype by blocking the formation of meiotic DSBs, likely reflects the premeiotic downregulation of TORC1 activity by the GATOR1 complex in early oogenesis (Wei et al. 2014).

### GATOR1 and TSC promote the repair of meiotic DSBs

Hyperactivation of TORC1 has been linked to defects in the DNA damage response in single celled and multicellular organisms (Begley, Rosenbach, Ideker, & Samson, 2004; Feng et al., 2007; Klermund, Bender, & Luke, 2014; Ma, Vassetzky, & Dokudovskaya, 2018; Pai et al., 2016; Xie et al., 2018). The observation that meiotic DSBs promote the GATOR1 dependent downregulation of TORC1 activity during *Drosophila* oogenesis, suggested the low TORC1 activity may be important to the regulation of meiotic DSB. In our previous work, we found that GATOR1 mutant ovaries had TORC1 activity levels approximately three times of those observed in wild-type ovaries (Cai et al., 2016; Wei et al., 2014). Here we demonstrate that GATOR1 mutant ovaries exhibit multiple phenotypes consistent with the misregulation of meiotic DSBs including, an increase in the steady state number of Mei-W68/Spo-11 induced DSBs, the retention of meiotic DSBs into later stages of oogenesis and the hyperactivation of p53. (Cai et al., 2016; Wei et al., 2016). Importantly, RNAi depletions of *Tsc1* phenocopied the GATOR1 ovarian defects. Thus, the misregulation of meiotic DSBs observed in GATOR1 mutant oocytes are due to high TORC1 activity and not to a TORC1 independent function of the GATOR1 complex.

Epistasis analysis between the GATOR1 component *nprl3* and the Rad51 homolog *spnA*, strongly suggest that GATOR1 impacts the repair, rather than the generation, of meiotic DSBs. We determined that double mutants of *nprl2* and the Rad51 homolog *spnA*, which is required for the repair of meiotic DSBs, have approximately the same number of DSBs as *spnA* single mutants. These data are consistent with GATOR1 and *spnA* influencing the common process of DNA repair and are inconsistent with GATOR1 mutants producing supernumerary breaks.

Our observations on the role of the GATOR1 complex during *Drosophila* oogenesis are particularly intriguing in light of similar meiotic defects observed in a *npr3* mutants in *Saccharomyces cerevisiae* (Jordan et al., 2007). In the sporulation proficient strain SK1, *npr3* mutant cells enter meiosis and express the transcription factor and master regulator of gametogenesis IME1 with wild-type kinetics (Jordan et al., 2007). Subsequently, *npr3* mutants exhibit a mild delay in the generation of meiotic DNA breaks, but a substantial delay in the repair of meiotic DSBs (Jordan et al., 2007). Thus, yeast and *Drosophila* SEACIT/GATOR1 mutants share a common meiotic phenotype, the delayed repair of meiotic DSBs. These results raise the intriguing possibility that low TORC1 activity may be a common feature of the early meiotic cycle in many organisms.

### GATOR1 opposes retrotransposon expression

The initiation of homologous recombination through the programmed generation of DNA double-stranded breaks (DSBs) is a universal feature of meiosis (Gray & Cohen, 2016; McKim & Hayashi-Hagihara, 1998). DSBs represent a dangerous form of DNA damage that can result in dramatic and permanent changes to the germline genome (Alexander et al., 2010). To minimize this destructive potential, the generation and repair of meiotic DSBs is tightly controlled in space and time (Longhese, Bonetti, Guerini, Manfrini, & Clerici, 2009). The activation of transposable elements represents an additional threat to genome integrity in germ line cells (Crichton, Dunican, Maclennan, Meehan, & Adams, 2014; Toth, Pezic, Stuwe, & Webster, 2016). Genotoxic stress, resulting from DNA damage, has been implicated in the deregulation of transposons in multiple organisms (Beauregard, Curcio, & Belfort, 2008; Bradshaw & McEntee, 1989; Hagan, Sheffield, & Rudin, 2003; Walbot, 1992). Thus, germ line cells may be at an increased risk for transposon derepression due to the genotoxic stress associated with meiotic recombination. Consistent with this hypothesis, germ line cells have evolved extensive surveillance systems to detect and silence transposons beyond the surveillance systems present in most somatic tissues (Khurana & Theurkauf, 2010; Ku & Lin, 2014; Toth et al., 2016).

Genotoxic stress has been implicated in the deregulation of retrotransposon expression or activation in multiple organisms including *Drosophila* (Beauregard et al., 2008; Bradshaw & McEntee, 1989; Hagan et al., 2003; McClintock, 1984; Molla-Herman, Valles, Ganem-Elbaz, Antoniewski, & Huynh, 2015; Walbot, 1992; Wylie et al., 2016). In line with these studies, we find that in GATOR1 mutants, the DSBs that initiate meiotic recombination trigger the deregulation of retrotransposon expression. Similarly, *p53* mutant females derepress retrotransposon expression during oogenesis, but as observed in GATOR1 mutants, primarily in the presence of meiotic DSBs (Wylie et al., 2016). Double mutants of *nprl3, p53* exhibit a dramatic increase in retrotransposon expression relative to either *p53* or *nprl3* single mutants, implying that *p53* and GATOR1 act through independent pathways to repress retrotransposon expression in the female germline.

Our data suggest that the GATOR1 complex may influence retrotransposon expression independent of the regulation of TORC1 activity. While both GATOR1 and TSC are required for the efficient repair of meiotic DSBs, in contrast to GATOR1 mutant ovaries, we observed little to no increase in retrotransposon expression in the *Tsc1* depleted ovaries. While this may reflect the incomplete depletion of *Tsc1*, a second possibility is that the GATOR1 complex inhibits retrotransposon expression independent of TORC1 inhibition. Intriguingly, GATOR1 components, but not TSC components, were recently identified in a high throughput screen for genes that suppress (Long Interspersed Element-1) LINE1 expression in mammalian tissue culture cells (N. Liu et al., 2018). Taken together, these data hint that the GATOR1 complex may impact retrotransposon expression in the germline via two independent pathways: First by promoting the repair of meiotic DSBs through the downregulation of TORC1 activity and second via a pathway that functions independent of TORC1 inhibition.

Genes encoding components of the GATOR1 complex are often deleted in cancers (Bar-Peled et al., 2013; Ji et al., 2002; Lerman & Minna, 2000; Ueda et al., 2006). As is observed in GATOR1 mutants, cancer cells frequently have increased TORC1 activity, increased genomic instability and increased retrotransposon expression. Thus, in the future it will be important to identify the molecular mechanism by which the GATOR1 complex influences both the response to genotoxic stress and the expression of retrotransposons under both normal and pathological conditions.

## Materials and methods

### Fly stocks

All fly stocks were maintained at 25°C on standard media. The *p53R-GFP* transgenic line was a gift from John M. Abrams (Lu et al., 2010). The germline specific driver nanos-Gal4 was obtained from Ruth Lehmann (Van Doren, Williamson, & Lehmann, 1998). The *spnA*^*093A*^stock was a gift from Ruth Lehmann (Staeva-Vieira et al., 2003). The *nprl2*^*1*^, *nprl3*^*1*^, *iml1*^*1*^, and *UAS-Nprl3* were described previously (Cai et al., 2016; Wei et al., 2016). The stocks *w*^*1118*^*; Df(3L)ED4515, P{3’.RS5+3.3’}ED4515/TM6C, cu*^*1*^ *Sb*^*1*^ (BDSC#9071), *w*^*1118*^*; Df(3L)ED4238, P{3’.RS5+3.3’}ED4238/TM6C, cu*^*1*^ *Sb*^*1*^ (BDSC#8052), *w*^*1118*^*; P{neoFRT}82B P{Ubi-mRFP.nls}3R* (BDSC#30555), *P{hsFLP}22, y*^*1*^ *w*^*\**^*; P{arm-lacZ.V}70C P{neoFRT}80B* (BDSC#6341), *MTD-GAL4 (P{w[+mC]=otu-GAL4::VP16.R}1, w[*] P{w[+mC]=GAL4-nos.NGT}40; P{w[+mC]=GAL4::VP16 nos.UTR}CG6325[MVD1]*, BDSC#31777), UAS-Tsc1 RNAi (y^1^ sc^*^ v^1^; P{TRiP.GL00012}attP2, BDSC#35144), UAS-mCherry RNAi (y^1^ sc^*^ v^1^; P{VALIUM20-mCherry}attP2), *y*^*1*^ *w*^*1118*^*; p53*^*5A-1-4*^ (BDSC#6815), *y*^*1*^ *w*^*1*^*/Dp(1;Y)y*^*+*^*; mei-P22*^*P22*^; *sv*^*spa-pol*^ (BDSC#4931), *Dp(1;Y)B*^*S*^*; ru*^*1*^ *st*^*1*^ *e*^*1*^ *spn-A*^*1*^ *ca*^*1*^*/<u>TM3</u>, Sb*^*1*^ (BDSC#3322) were obtained from Bloomington Stock Center.

### Western blot analysis

The protocol was adapted from Bjedov I (2010) (Bjedov et al., 2010). Briefly 6 pairs of ovaries were freshly dissected in cell insect media and homogenized in 30µl of 4x Laemmli loading sample buffer (Invitrogen, #NP0008) containing 10x sample reducing agent (Invitrogen, #NP009). Extracts were cleared by centrifugation and boiled for 10 minutes at 90°C. 10µl of protein extract was loaded per lane on polyacrylamide gel (Invitrogen, #NP0335). Proteins were separated and transferred to nitrocellulose membrane. Primary antibodies used were as follows: guinea pig anti-dS6K (gift of Aurelius Teleman,1:5,000,) (Hahn et al., 2010) and rabbit anti-phospho-Thr398-S6K (Cell Signaling Technologies #9209, 1:1,000). HRP-conjugated secondary antibodies (Jackson Immunoresearch, AffiniPure anti-rabbit #111-005-144 and anti-guinea pig #106-005-003) were used. Blots were developed using the ECL detection system (PerkinElmer, #NEL105001EA). Western blots were analyzed using ImageJ program (US National Institutes of Health).

### Immunofluorescence and microscopy

Immunofluorescence was performed as previously described (Hong et al., 2003; Iida and Lilly, 2004). Primary antibodies used were as follows: rabbit anti-GFP (Invitrogen, 1:1000); rabbit anti-γ– H2Av (Active Motif, 1:500); rabbit anti-C(3)G 1-3000 (Hong, et. al., 2003); mouse anti-1B1 (Developmental Studies Hybridoma Bank, 1:100); mouse anti-γ-H2Av (Developmental Studies Hybridoma Bank, 1:5000); mouse anti-C(3)G (kindly provided by R. Scott Hawley, 1:200) (Page & Hawley, 2001). Alexa-488 and Alex-594 (Invitrogen, 1:1000) secondary antibodies were used for fluorescence. After staining, ovaries were mounted in prolong gold antifade reagent with DAPI (Life Technology). Images were acquired on either a Leica SP5 confocal microscope or Zeiss LSM 880 with Airyscan confocal microscope.

### γ-H2Av foci quantification

To score the number of γ-H2Av foci per oocyte, ovaries were stained with antibodies against C(3)G and γ-H2Av as well as the DNA dye DAPI. Pro-oocytes and oocytes were identified by the pattern of anti-C(3)G staining (Page and Hawley 2001). Multiple Z-sections encompassing an entire region 2a pro-oocyte nucleus were acquired by Leica SP5 confocal microscope or Zeiss LSM 880 with Airyscan confocal microscopy. The obtained z-stacks of images were deconvolved to remove out-of-focus light and z-distortion with Huygens Professional software (Scientific Volume Imaging) and clearly defined γ-H2Av were counted manually. Alternatively, 3D images were rendered by using Imaris software (Bitplane) and a graph workstation equipped with NVIDIA Quadro 3D vision system. Clearly defined γ-H2Av foci were visualized and counted by using Imaris spots module or image J to define the total number of foci per nucleus.

### MMS sensitivity assay

The assay was performed as described in Ghabrial and Schupbach (1998). Briefly, 10 males and 10 virgin females were mated in vials for 2 days at 25°C. Parents were transferred into German food (Genesee Scientific, Cat#66-115, Day 1) for 24 hours at 25°C and allowed to lay eggs. On Day 2 the parents were removed and eggs were allowed to mature for 24 hours. Subsequently, the first and second instar larvae were treated with 250 µL of either 0.04% or 0.08% of the mutagen methyl methanesulfonate (MMS) (Sigma, Cat#129925-5G). Control larvae were treated with 250 µL water. After eclosion, the number of heterozygous and homozygous mutant flies were determined, and the ratio of each genotype was calculated and divided by expected ratios.

### Retrotransposon Expression Analysis

Total RNA was isolated from dissected ovaries using the RNeaszy Kit (Qiagen) and treated with DNase. cDNA was generated using High-capacity cDNA Reverse Transcription Kit (Thermo Fisher). Real-time PCR was performed with Power SYBR green Mastermix (Thermo Fisher) using the following primers.

Rp49 Forward: CCGCTTCAAGGGACAGTATC;

Rp49 Reverse: GACAATCTCCTTGCGCTTCT;

TAHRE Forward: CTGTTGCACAAAGCCAAGAA;

TAHRE Reverse: GTTGGTAATGTTCGCGTCCT;

Het-A Forward: TCCAACTTTGTAACTCCCAGC;

Het-A Reverse: TTCTGGCTTTGGATTCCTCG;

Idefix Forward: TGAAGAAAAGAAGGGCGAGA;

Idefix Reverse: TTCTGCTGTTGATGCTTTGG;

Gypsy Forward: CCAGGTCGGGCTGTTATAGG;

Gypsy Reverse: GAACCGGTGTACTCAAGAGC.

The rp49 was used for normalization.

## Acknowledgements

Multiple stocks used in this study were obtained from the Bloomington *Drosophila* Stock Center (supported by NIH Grant P40OD018537). Research reported in this publication was supported by the *Eunice Kennedy Shriver* National Institute of Child Health and Human Development Intramural Research program (to M.A.L., HD00163 16).

